# Computational reconstruction of Tyrannosaurus rex collagen for biomanufacturing of paleo-inspired biomaterials with tuneable stiffness

**DOI:** 10.64898/2025.12.15.689182

**Authors:** T Mitchell, E Wolvetang

## Abstract

Ancient protein fragments preserved in fossilized tissues offer evolutionary sequence information inaccessible through conventional genomics. Here we demonstrate computational reconstruction of full-length Tyrannosaurus rex type I collagen (COL1A1) by integrating paleo-proteomics data with protein language models. Using TOC large protein model with pattern-constrained generation to maintain collagen’s Gly-X-Y triplet architecture, we generated five computationally designed sequences incorporating validated T. rex peptide fragments. These generated variants maintain triple helix structural features (23.0% Gly-X-Y content) and exhibit enhanced lysine content (up to 84 residues vs. 57 in wild-type chicken) to increase enzymatic crosslinking capacity. To enable spatial and temporal control of material properties such as stiffness, we engineered a doxycycline-inducible lysyl oxidase-4 (LOX4) expression system in TOC t-rex cell line that achieved 9-fold enzyme induction. This biomimetic approach, inspired by functionally graded fish scale architecture, decouples constitutive collagen production from inducible crosslinking, enabling biomanufacturing of materials with tuneable mechanical gradients. We validate stable transgene integration, confirm tight transcriptional control, and demonstrate system compatibility with scalable cell culture. This work establishes a platform combining paleo-proteomics, protein language modelling, and bioinspired engineering to produce next-generation biomaterials from computationally reconstructed extinct proteins.

## Introduction

Protein language models (PLMs) trained on evolutionary sequence databases can generate novel functional proteins across diverse families ^1–3^, yet their application to structurally constrained proteins like fibrillar collagens remains challenging ^4,5^. Type I collagen, the most abundant structural protein in vertebrates, requires strict Gly-X-Y repeat periodicity for triple helix formation, a constraint not reliably learned by current generative models ^6^. Meanwhile, discovery of endogenous collagen peptides in 68-million-year-old Tyrannosaurus rex fossils provides evolutionary sequence data that, while fragmentary, offers phylogenetically distant variants potentially encoding collagen species that may impart distinct material properties to end-products made from cells expressing such novel collagens ^7,8^.

Collagen mechanical performance depends critically on enzymatic crosslinking through lysyl oxidase (LOX) enzymes, which catalyse oxidative deamination of lysine/hydroxylysine to form covalent hydroxylysyl-pyridinoline (HP) crosslinks ^9,10^. Natural systems achieve functionally graded materials through spatial control of LOX expression gene copies, thus enabling tissue-specific crosslink patterns. For example, piranha-resistant Arapaima gigas scales combine heavily crosslinked hard outer layers with flexible collagenous cores for exceptional impact resistance ^11,12^.

Here we integrate paleo-proteomics, constrained protein language modelling, and biomimetic engineering to: (1) computationally reconstruct a full-length T. rex-inspired COL1A1 using phylogenetically informed pattern-constrained generation; (2) characterize sequences for collagen-relevant structural features and crosslinking potential; (3) engineer an inducible LOX4 system enabling spatial control of collagen crosslinking; and (4) validate system functionality through stable cell line generation and transcriptional profiling. This approach bridges computational protein design with biomanufacturing to produce functionally graded biomaterials that should mimic natural dermal armour architectures.

## Results

### Pattern-constrained generation maintains collagen architecture while incorporating ancient sequences

Unconstrained sequence generation using TOC’s large protein model failed to preserve the Gly-X-Y repeat pattern essential for collagen triple helix formation, yielding sequences with <2% valid triplet content versus 21.9% in wild-type chicken COL1A1. This failure demonstrates that large protein models, despite training on >1 billion protein sequences, does not reliably learn the strict architectural constraints of fibrillar collagens.

To overcome this limitation, we implemented pattern-constrained generation. Using chicken COL1A1 (UniProt P02457, 1,453 amino acids) as scaffold based on phylogenetic proximity to T. rex fragments ^7^, we: (1) locked all glycine residues at position 1 of Gly-X-Y triplets (494 positions, 34.0% of sequence); (2) inserted and locked validated T. rex fragments; and (3) enabled the TOC large protein model to sample at 957 modifiable X and Y positions (66.0% of sequence) using temperature T=0.8, top-p=0.9 nucleus sampling.

Five independent sequences (Gen4-1 through Gen4-5, 1451 amino acids each) were generated with random seeds 42-46. All sequences maintained 23.0% Gly-X-Y content, closely matching wild-type chicken COL1A1 (21.9%) and confirming successful preservation of triple helix architecture (Fig. 1a). Proline analysis revealed 270-320 total proline residues per sequence with 50-75 in Y-positions. This represents substantial hydroxyproline potential (17-24% of sequence) exceeding wild-type chicken’s 47 Y-position prolines (Fig. 1b). Key triplet motif quantification showed enrichment of GPK motifs (Gly-Pro-Lys, 9-14 occurrences) critical for enzymatic crosslinking, with Gen4-3 containing 14 GPK sites versus 9 in wild-type (Fig. 1c). Gen4-3 selected as lead candidate based on maximum lysine content (84 residues) for enhanced crosslinking capacity.

**Figure 1:**
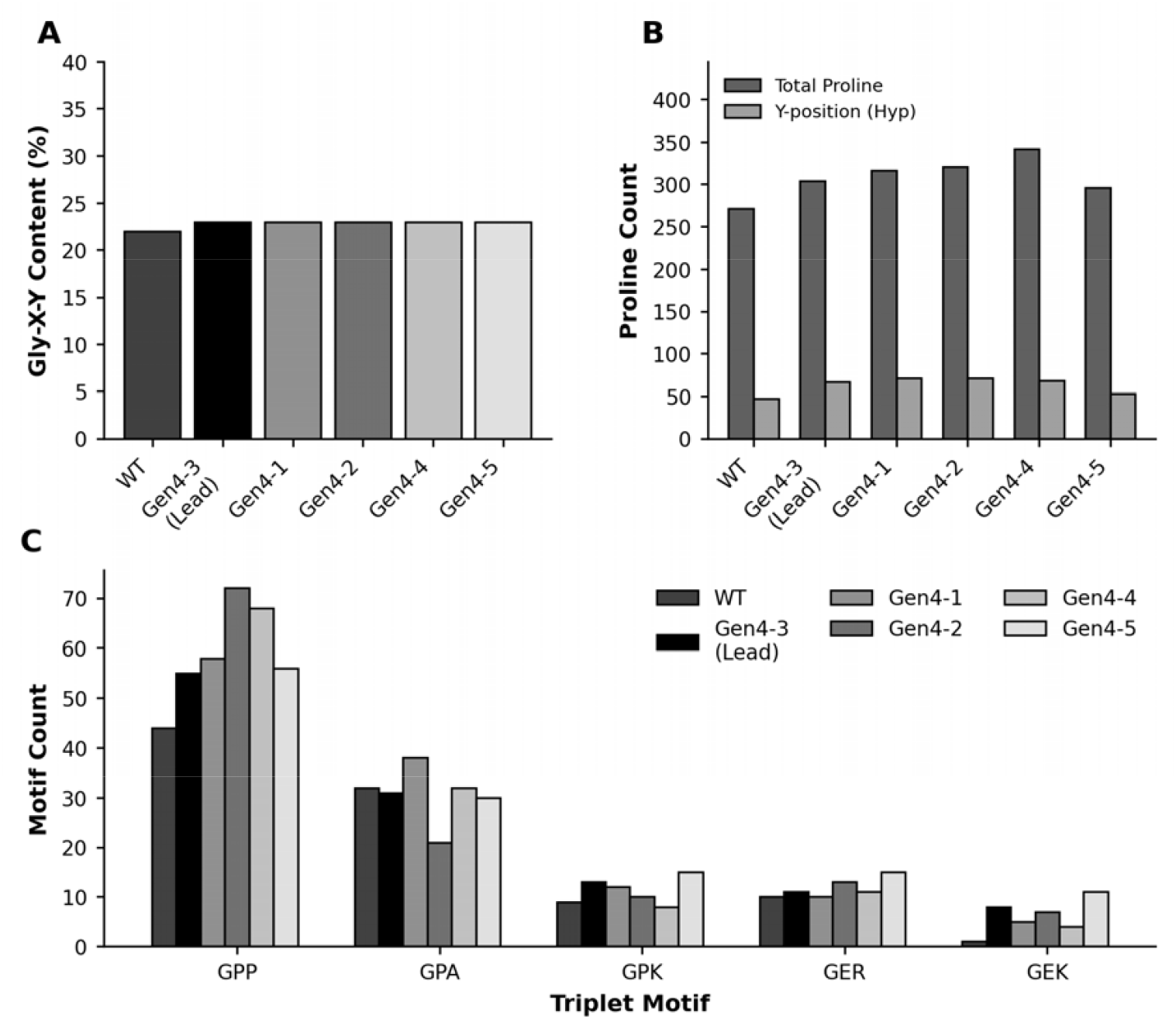
Computational design strategy for lysine-enriched T. rex collagen variants. (A)Gly-X-Y triplet content (%) in wild-type (WT) chicken COL1A1 and five computationally designed T. rex collagen variants (Gen4-1 through Gen4-5). Gen4-3 was selected as the lead candidate. (B) Total proline count and Y-position proline count (hydroxyproline potential) across all sequences. (C) Frequency of key collagen triplet motifs (GPP, GPA, GPK, GER, GEK) demonstrating retention of critical structural elements in designed sequences.

### Generated sequences exhibit enhanced lysine content with preferential Y-position enrichment

Amino acid composition analysis revealed that generated sequences preserve collagen biochemical signatures, high glycine (26.9 ± 0.2%) and proline (19.9 ± 1.0%), while exhibiting systematic lysine enrichment (Fig. 2). Wild-type chicken COL1A1 contains 57 lysines (3.9%), whereas generated sequences ranged from 60-84 lysines (4.1-5.8%), with Gen4-3 showing maximum enrichment (47% increase).

**Figure 2:**
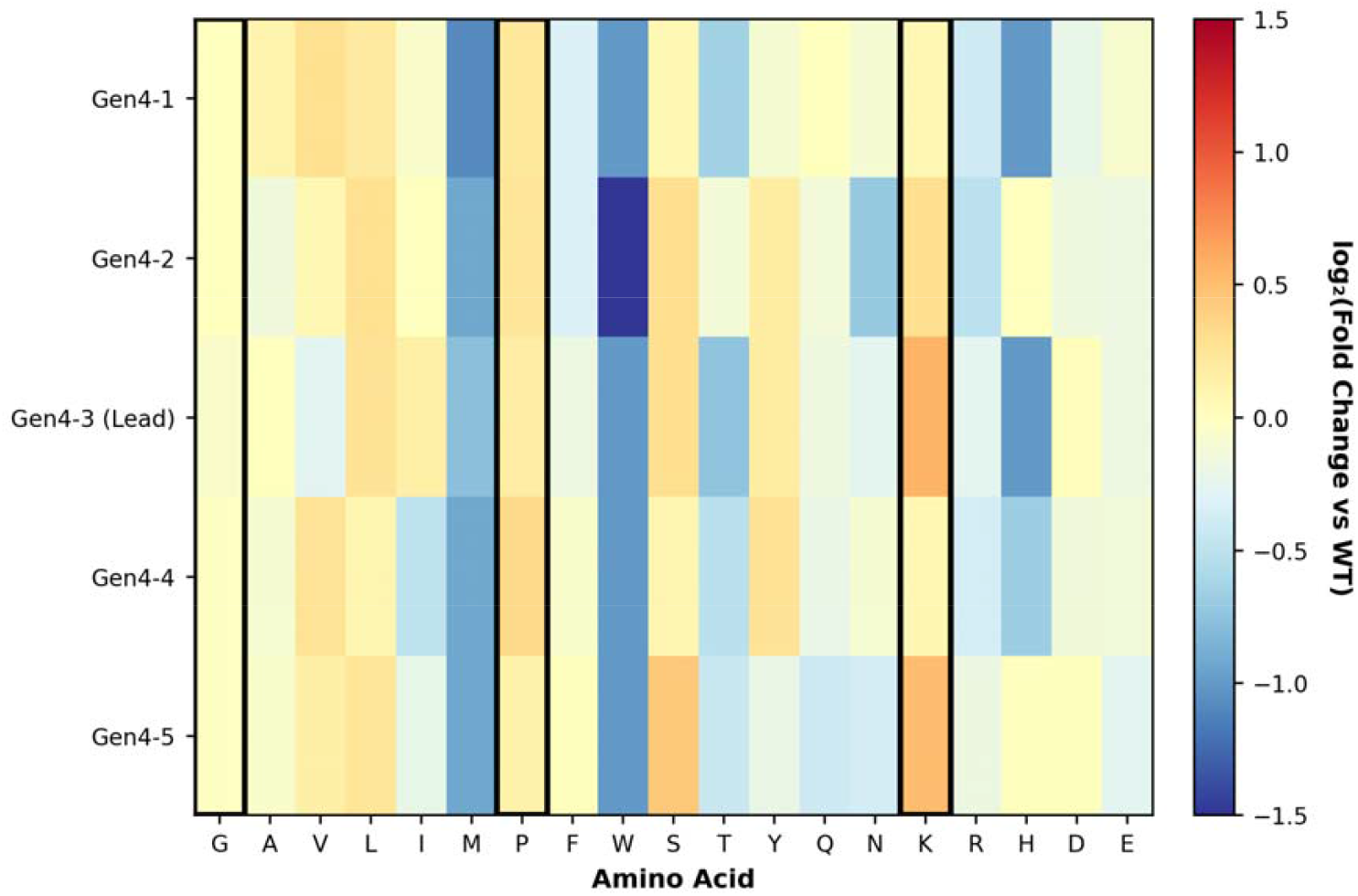
Amino acid composition comparison of designed sequences. Heatmap depicting log2 fold-change in amino acid composition for all five generated sequences (Gen4-1 through Gen4-5) relative to WT chicken COL1A1. Black rectangles highlight key structural residues (Gly, Pro, Lys), showing targeted lysine enrichment while maintaining collagen-characteristic composition.

To quantify structural differences between generated sequences, we performed principal component analysis on six biophysical features (Gly-X-Y content, amino acid composition, instability, GRAVY) (Fig. 3a). PC1 (43.5% variance) was dominated by proline content (0.999 loading), instability index (0.973), and Gly-X-Y content (0.955), separating sequences by their collagen-characteristic structural features (Fig. 3b). PC2 (37.8% variance) showed strongest loadings for glycine percentage (1.027) and lysine percentage (-0.916), reflecting amino acid composition relevant to crosslinking capacity (Fig. 3c).

**Figure 3:**
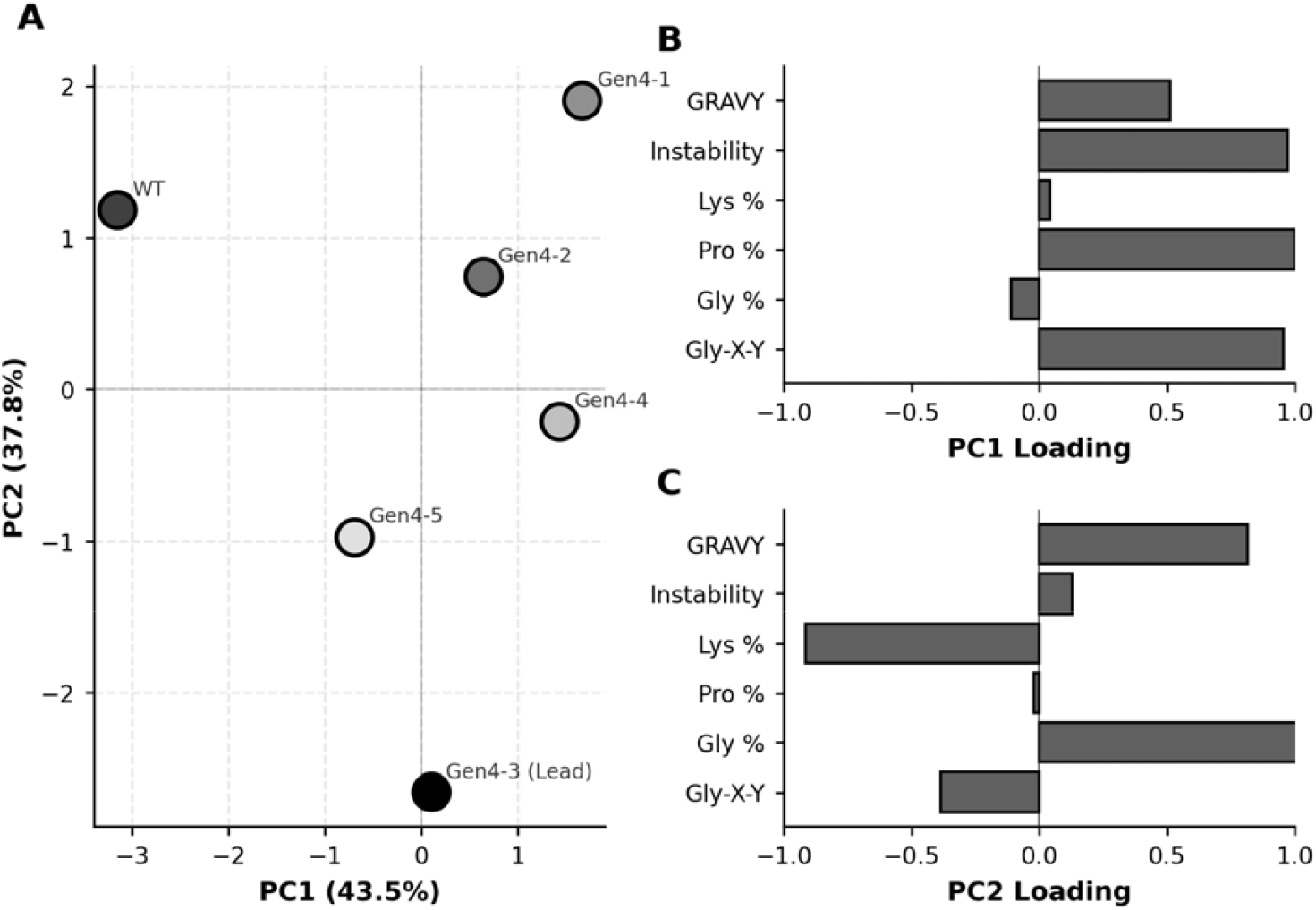
Principal component analysis of structural features. (A)PCA of six structural features (Gly-X-Y content, glycine %, proline %, lysine %, instability index, GRAVY score) showing relationships among WT and all five designed variants. Gen4-3 (Lead) demonstrates optimal balance of collagen-like properties and enhanced lysine content. (B) Feature loadings for PC1 (explaining primary axis of variation). (C) Feature loadings for PC2 (explaining secondary axis of variation).

Wild-type chicken COL1A1 occupied a distinct position (PC1 = -3.15, PC2 = 1.19), while generated sequences distributed broadly across both axes. Gen4-3 (Lead) showed the most extreme divergence from wild-type (PC1 = 0.10, PC2 = -2.65), driven primarily by its exceptionally high lysine content (84 residues, 5.8%), which contributes to the strong negative PC2 score. This combination of moderate collagen structure (PC1 near zero) with maximal crosslinking substrate availability (negative PC2) makes Gen4-3 optimal for biomimetic materials requiring enhanced mechanical properties through enzymatic crosslinking.

Detailed lysine distribution analysis revealed preferential positioning in Y locations of Gly-X-Y triplets, where they serve as substrates for lysyl hydroxylase to generate hydroxylysine, the preferred substrate for LOX-mediated crosslink formation^9,13^. Gen4-3 contains 25 Y-position lysines versus 9 in wild-type (2.8-fold increase) (Fig. 4a,b). Sliding window analysis (30 amino acid window, 10 amino acid step) showed regional lysine density peaks exceeding 13% local concentration in Gen4-3, compared to 6-8% maximum in wild-type (Fig. 4c). This non-uniform distribution creates potential “hotspots” for spatially controlled crosslinking.

**Figure 4:**
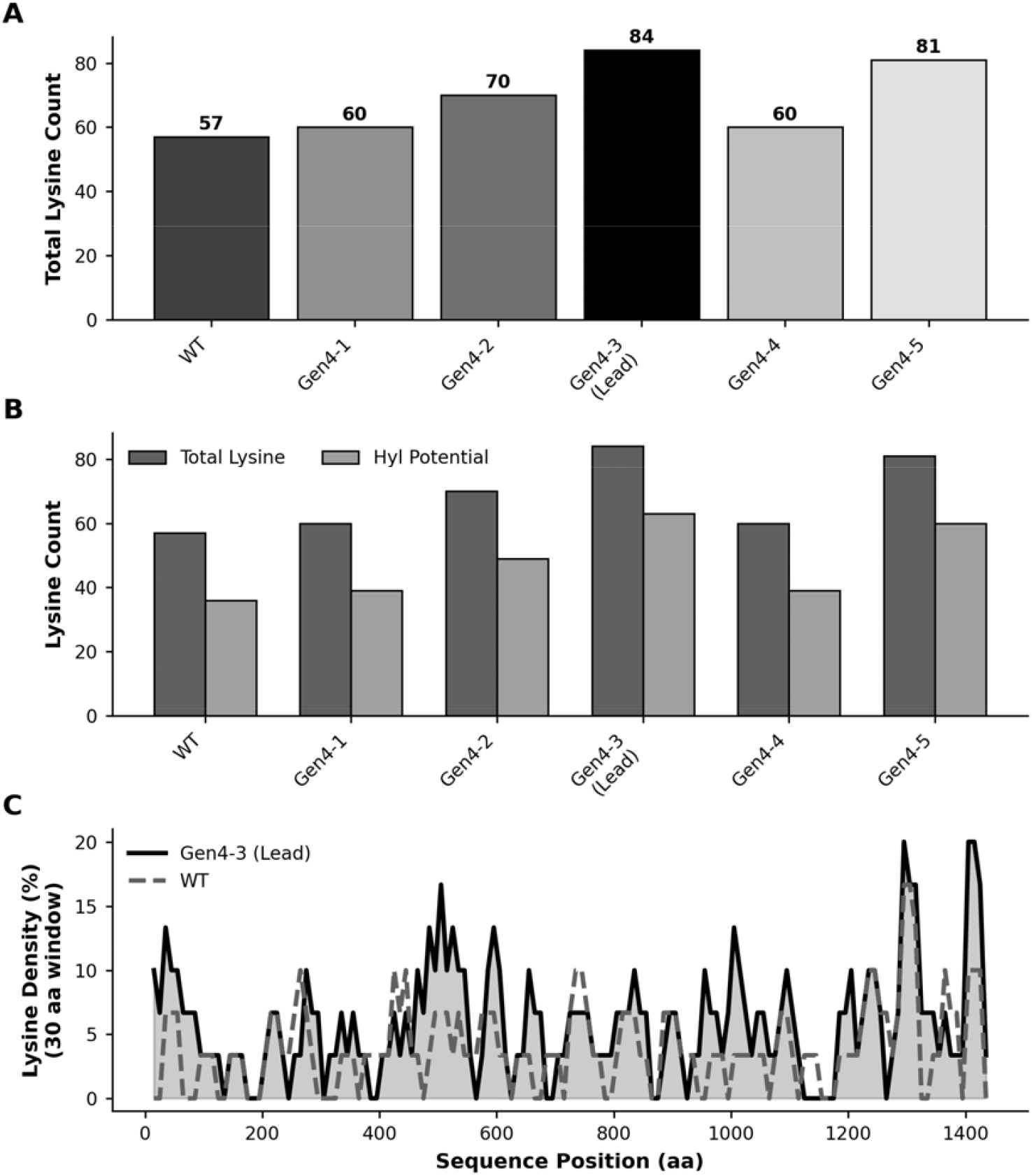
Lysine enrichment and distribution in designed sequences. (A) Total lysine count per sequence showing significant enrichment in designed variants versus WT. (B) Total lysine versus lysine in X/Y positions (hydroxylysine potential), demonstrating strategic placement for post-translational modification. (C) Sliding window analysis (30 amino acid window, 10 amino acid step) of lysine density comparing Gen4-3 (Lead) to WT chicken COL1A1, revealing engineered high-density lysine regions for enhanced crosslinking capacity.

### Biomimetic expression system enables decoupled collagen production and crosslinking control

To translate computational designs into bio-manufacturable materials with tuneable properties, we engineered a dual-cassette expression system enabling spatial and temporal control of collagen crosslinking (Fig. 5a). The PiggyBac transposon construct contains: (1) CMV promoter driving constitutive expression of Gen4-3 COL1A1-FLAG fusion protein for continuous collagen substrate production; and (2) tetracycline response element (TRE) promoter driving doxycycline-inducible LOX4 (UniProt A0A8V0Z6H4, Atlantic salmon) for controlled crosslink formation. This architecture decouples substrate accumulation from enzymatic modification, enabling: (i) pre-accumulation of un-crosslinked collagen matrix; (ii) precise temporal initiation of crosslinking; and (iii) spatial patterning through localized doxycycline application to recreate functionally graded architectures found in natural dermal armour^12^.

**Figure 5:**
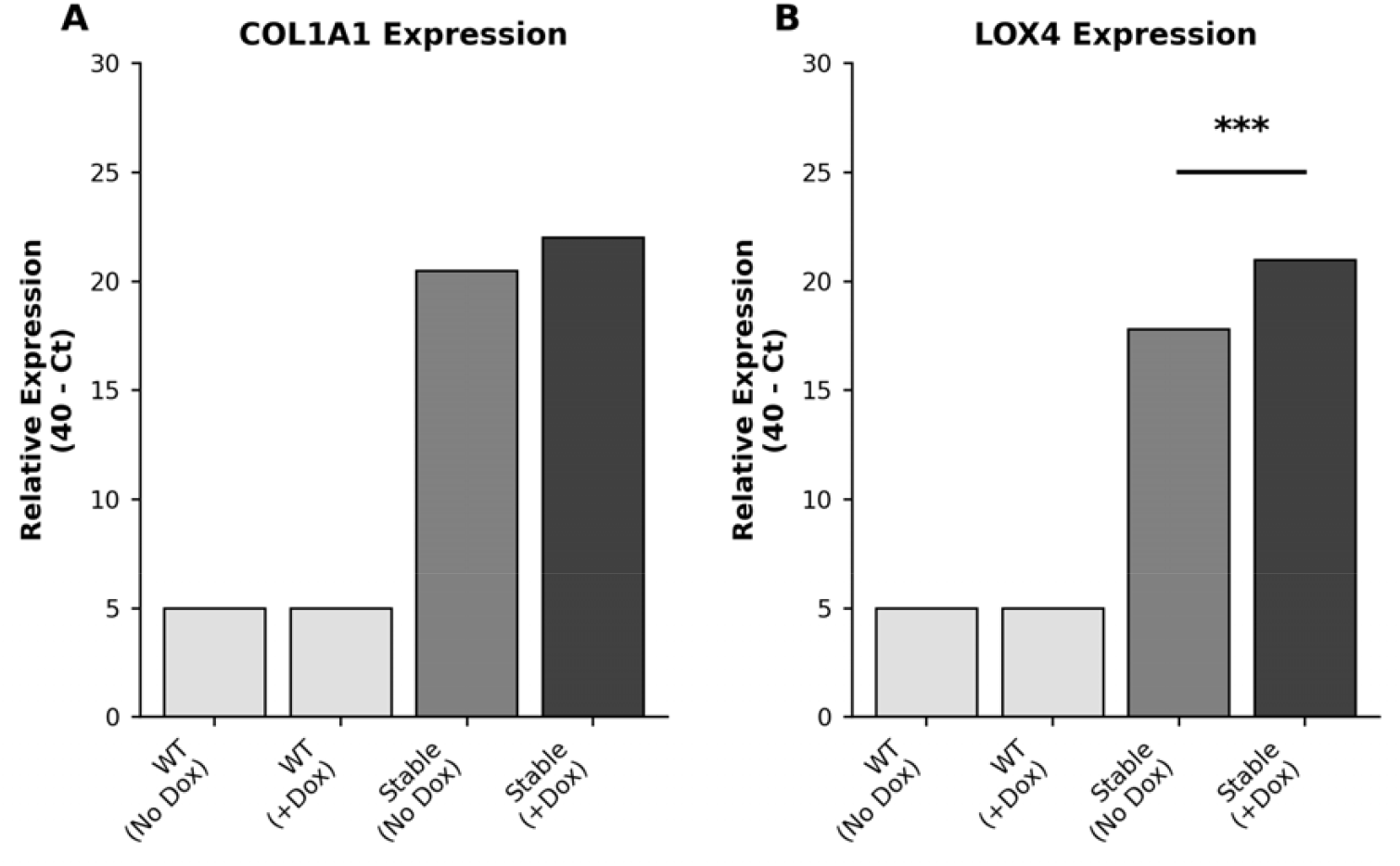
Experimental validation of transgene expression in mammalian cells. (A) Expression of chicken COL1A1 transgene (Gen4-3 lead variant) in WT untransfected cells (negative control) and stable transfected cell lines, measured by qRT-PCR with and without doxycycline (Dox) induction (n=3 biological replicates). WT cells show no expression (Ct ∼35, background). (B) Expression of LOX4 (encoding lysyl hydroxylase, collagen crosslinking enzyme) showing significant induction upon Dox treatment in stable cells (***P < 0.001, two-tailed Student’s t-test; n=3 biological replicates). Data in (A) and (B) are presented as relative expression (40 - Ct) ± s.e.m.

LOX4 was selected over other LOX family members based on substrate specificity for hydroxylysine and high activity toward HP crosslink formation^13^. The constitutive/inducible architecture mimics natural systems where collagen synthesis and crosslinking are temporally and spatially regulated during tissue development.

### Stable integration and transcriptional profiling validate system functionality

Co-transfection of TOC t-rex cell line with three PiggyBac plasmids (1) dual-cassette COL1A1-FLAG/TRE-LOX4 construct, (2) rtTA tetracycline trans-activator, and (3) hyperactive PiggyBac transposase followed by dual antibiotic selection (puromycin 2.0 μg/mL, hygromycin B 400 μg/mL) yielded a polyclonal stable cell line. Morphological assessment confirmed normal cell viability under both uninduced and induced (2 μg/mL doxycycline, 48 hours) conditions, indicating that high-level expression of engineered T. rex collagen and LOX4 is well-tolerated.

Quantitative RT-PCR with GAPDH normalization revealed constitutive COL1A1 expression in stable cells (Ct = 19.54 ± 0.06 uninduced, Ct = 18.03 ± 0.06 induced), while wild-type TOC cell line showed no detectable expression (Ct undetermined). Doxycycline treatment resulted in 2.92-fold COL1A1 induction (ΔΔCt = 2.92), potentially reflecting doxycycline-mediated transcriptional enhancement or mRNA stabilization effects in the master cell line (Fig. 5a). Critically, basal COL1A1 expression (Ct ∼19.5) confirms substrate availability prior to LOX4 induction, as required for the decoupled production/crosslinking strategy.

LOX4 expression demonstrated tight inducible control: uninduced cells showed low basal expression (Ct = 22.21 ± 0.04), while doxycycline treatment induced robust 9.09-fold upregulation (ΔΔCt = 9.09, ΔΔCt = -3.18, induced Ct = 19.06 ± 0.02) (Fig. 5b). This ΔCt shift of 3.2 cycles, corresponding to ∼9 fold higher LOX4 mRNA, should provide sufficient dynamic range to create spatial crosslinking gradients, as seen in teleost tissue^14^.

### System enables biomanufacturing of functionally graded materials

The validated expression system enables a biomanufacturing workflow for producing functionally graded collagenous biomaterials: (1) stable cell line expansion in standard culture conditions to accumulate cell mass; (2) three-dimensional tissue culture (scaffold-based or scaffold-free) to deposit collagen extracellular matrix; (3) spatial patterning of LOX4 induction through layered doxycycline application, high concentration in outer regions for heavily crosslinked hard layers, low/no induction in core regions for flexible layers; and (4) decellularization and processing to yield acellular collagenous material with designed mechanical gradients.

This approach directly mimics the hard exterior/flexible interior architecture of Arapaima gigas scales, which achieve exceptional puncture resistance through graded crosslink density ^12^: mineralized, heavily crosslinked outer layers provide hardness to resist initial impact, while softer inner layers absorb energy and prevent crack propagation. By controlling local doxycycline concentration during tissue culture, we can recreate this evolutionary strategy in bio-manufactured materials using computationally designed T. rex collagen optimized for enzymatic crosslinking (84 lysines vs. 57 in wild-type chicken).

Spatially localized doxycycline-soaked beads would provide a practical way to program these gradients, generating leather-like materials with deliberately heterogeneous surface patterns and region-specific mechanical properties that are not achievable with traditional, uniformly tanned hides.

Material characterization experiments will assess: (1) hydroxylysyl-pyridinoline (HP) crosslink density as a function of LOX4 induction level; (2) mechanical properties across regions with differential crosslinking; and (3) correlation between LOX4 expression level, HP crosslink density, and resulting mechanical performance.

## Discussion

This work establishes a biotechnology platform integrating three key innovations: (1) computational reconstruction of extinct proteins using pattern-constrained protein language models; (2) rational design for enhanced functionality (lysine enrichment for crosslinking); and (3) biomimetic manufacturing systems enabling spatial control of post-translational modifications to create functionally graded materials.

### Pattern-constrained generation as a generalizable strategy for structurally complex proteins

The failure of unconstrained large protein models to maintain Gly-X-Y periodicity, contrasted with success of pattern-constrained generation, reveals limitations and opportunities for PLM-based protein design. While models trained on evolutionary sequence databases capture many functional constraints, strict architectural requirements, especially those involving repetitive structural motifs (collagen Gly-X-Y, silk fibroin repeats, elastin VPGXG pentapeptides), may require explicit enforcement during generation.

Our pattern-constrained approach represents a hybrid strategy: leveraging PLM knowledge of evolutionary sequence space for positions where diversity is tolerated, while enforcing structural constraints at positions critical for fold integrity. This framework is generalizable to other repeat proteins and could accelerate design of biomaterials with defined mechanical properties. Importantly, the model’s convergent enrichment of proline in Y positions (66.6 ± 7.5 residues vs. 47 in wild-type), despite no explicit instruction, demonstrates that PLMs trained on eukaryotic proteomes implicitly learn that this position undergoes post-translational hydroxylation, a critical stabilizing modification^15,16^.

### Enhanced lysine content enables tuneable crosslinking density

Gen4-3’s 84 lysine residues (5.8% of sequence) represent a 47% increase over wild-type chicken COL1A1, with critical enrichment at Y positions where lysyl hydroxylase generates hydroxylysine, the preferred LOX substrate^13^. This positions Gen4-3 as an optimized substrate for enzymatic crosslinking without compromising triple helix stability (23.0% Gly-X-Y content maintained). The non-uniform spatial distribution of lysine-rich regions creates potential “hotspots” that could concentrate crosslinks, potentially improving mechanical reinforcement through localized rather than uniform crosslinking.

Natural systems demonstrate the functional importance of crosslink density control: teleost fish scales with higher HP crosslink content show increased tensile strength and puncture resistance^12^, while mammalian tendons exhibit tissue-specific crosslink patterns correlating with mechanical demands^17^. Our 9-fold LOX4 induction range provides sufficient control to recreate such natural gradients in bio-manufactured materials.

### Translational potential for sustainable biomaterials

The environmental and ethical concerns of traditional leather production, including chromium tanning waste, animal welfare, and land use^18^, have driven interest in cell-cultured alternatives. Existing approaches focus on bovine collagen, offering sustainability improvements but limited material innovation. By computationally accessing extinct species diverged by ∼68 million years of evolution, we introduce sequence variants potentially encoding distinct material properties while maintaining a sustainable, animal-free production platform.

T. rex as an apex predator likely possessed collagen variants optimized for robust dermal protection. While we cannot directly validate this hypothesis, the enhanced lysine content (resulting from PLM generation on diverse vertebrate sequences including species under high mechanical demands) and integration with biomimetic crosslinking control positions this system for producing high-performance biomaterials. Target applications include:

1. High-performance leather alternatives for automotive, fashion, and upholstery applications requiring puncture resistance and durability
2. Functionally graded biomedical materials combining stiff regions for structural support with compliant regions for tissue interfacing
3. Tissue engineering scaffolds with spatially controlled mechanical properties to guide cell differentiation and tissue morphogenesis
4. Protective coatings and films leveraging graded crosslinking for impact-resistant yet flexible surface treatments

Commercial viability depends on demonstrating superior performance-to-cost ratios compared to traditional and cell-cultured bovine leather. The stable cell line platform enables scalable bioreactor-based production, potentially achieving cost competitiveness within 5-10 years as cell culture manufacturing matures.

### Experimental validation confirms proof-of-concept

Our RT-qPCR results validate three critical design elements: (1) stable chromosomal integration of large constructs (>10 kb total) using PiggyBac transposition; (2) constitutive production of computationally designed T. rex collagen without cytotoxicity; and (3) tight, inducible LOX4 control with 9-fold dynamic range. The compatibility with TOC t-rex cell line is particularly advantageous, as our cell lines provide appropriate post-translational modification machinery (prolyl hydroxylase, lysyl hydroxylase) for collagen processing while avoiding regulatory concerns associated with mammalian cell culture^19^.

Next steps require protein-level validation: Western blotting with anti-FLAG antibodies to confirm COL1A1-FLAG protein production and secretion; LOX4 enzyme activity assays to verify functional crosslinking activity; immunofluorescence microscopy to assess extracellular matrix fibril assembly; and mass spectrometry to quantify post-translational modifications (hydroxyproline, hydroxylysine content).

Correlation between doxycycline concentration, LOX4 expression, HP crosslink density, and mechanical properties will establish dose-response relationships needed for manufacturing process control.

### Broader implications for protein engineering and paleo-proteomics

This work demonstrates that fragmentary protein sequence data from fossilized tissues, previously viewed primarily as evolutionary markers, can serve as constraints for generative protein design. As paleo-proteomics methods improve, expanding the library of ancient protein sequences^20^, this approach could access broader sequence space including extinct lineages adapted to extreme environments or unique ecological niches.

More broadly, our pattern-constrained generation framework addresses a current limitation of protein language models: their difficulty with structurally complex, repetitive proteins. As PLMs expand to larger parameter counts and multimodal training data (sequence + structure + function)^21,22^, integrating explicit structural constraints during generation may enable design of increasingly complex proteins including multi-domain enzymes, megadalton structural assemblies, and synthetic protein polymers.

## Conclusion

We demonstrate that integrating paleo-proteomics, pattern-constrained protein language modelling, and biomimetic engineering enables computational reconstruction of extinct structural proteins with enhanced functional properties. Generated T. rex collagen variants maintain critical architectural features while exhibiting 47% increased lysine content for enhanced crosslinking. Our validated expression system decouples constitutive collagen production from inducible LOX4-mediated crosslinking, enabling biomanufacturing of functionally graded materials inspired by natural dermal armour. This platform bridges computational protein design with biotechnology manufacturing to produce next-generation biomaterials from evolutionarily distant sequence variants, with applications spanning sustainable leather alternatives to tissue engineering scaffolds.

## Acknowledgments

TOC credits Bas Korsten, Global Chief Creative Officer, and VML with the concept and project formation to develop the T-Rex leather product.

## Supplemetary Material

Computational reconstruction of Tyrannosaurus rex collagen for biomanufacturing of paleo-inspired biomaterials with tuneable stiffness

## Materials and Methods

### Computational sequence reconstruction

#### T. rex peptide database search

Ancient collagen peptide fragments from *Tyrannosaurus rex* specimen MOR 1125 were used as constraints for sequence generation. Peptide sequences were searched against the UniProt database (release 2024_04) using BLAST to identify homologous collagen sequences across vertebrate species. Chicken COL1A1 (UniProt P02457, 1,451 amino acids) was selected as the reference template based on phylogenetic proximity to *Archosauria* and completeness of structural annotation.

#### Protein language model generation

We used a large protein model to generate full-length collagen variants. To preserve the essential Gly-X-Y triplet repeat architecture required for collagen triple helix formation, we implemented pattern-constrained generation where all glycine residues at triplet position 1 (494 positions, 34% of sequence) were locked to the template sequence. The *Tyrannosaurus rex* peptide fragments were inserted based on sequence alignment. Remaining positions were sampled from the model’s learned probability distributions to generate sequence diversity while maintaining phylogenetic plausibility. Five independent full-length sequences (Gen4-1 through Gen4-5) were generated and validated for Gly-X-Y content and fragment preservation.

#### Sequence analysis

Amino acid composition and biophysical features were calculated using BioPython (v1.81). Gly-X-Y triplet content was determined by sliding window analysis with frame size 3. Lysine distribution was analyzed by position within the Gly-X-Y framework (X, Y, or non-triplet positions) and by regional density using a 30-amino-acid sliding window with 10-amino-acid steps. Pairwise sequence identities were calculated using Needleman-Wunsch global alignment (BLOSUM62 substitution matrix) as implemented in BioPython. Principal component analysis was performed on six standardized biophysical features (Gly-X-Y content, glycine %, proline %, lysine %, instability index, GRAVY score) using scikit-learn (v1.3.2) with singular value decomposition.

### Plasmid construction and cell line generation

#### Expression vector design

A dual-expression construct was designed using the PiggyBac transposon system for chromosomal integration. The vector contains three expression cassettes: (i) a constitutive promoter driving the lead collagen variant (Gen4-3) with C-terminal epitope tag for immunodetection; (ii) a tetracycline-responsive element (TRE) promoter controlling lysyl oxidase 4 (LOX4, UniProt A0A8V0Z6H4) expression to enable inducible crosslinking; and (iii) a selection cassette for antibiotic resistance. The Gen4-3 sequence includes the native chicken COL1A1 signal peptide (positions 1-22) for secretion into the extracellular matrix. The complete construct was assembled using molecular cloning techniques and verified by Sanger sequencing.

#### Stable cell line generation

The TOC master cell line was maintained in high-glucose DMEM supplemented with 10% fetal bovine serum and antibiotics at 37°C with 5% CO□. Cells were co-transfected with the expression vector and PiggyBac transposase helper plasmid using a commercial lipid-based transfection reagent according to the manufacturer’s protocol. Transfected cells underwent dual antibiotic selection beginning 48 h post-transfection. Resistant colonies were expanded as pooled populations and screened for transgene expression by qRT-PCR. High-expressing stable pools were maintained under continuous selection and used for all subsequent experiments.

### Gene expression analysis

#### RNA extraction and qRT-PCR

Stable cell lines were cultured with or without doxycycline (2 μg/mL) for 48 h to induce LOX4 expression. Total RNA was extracted using column-based purification kits following the manufacturer’s instructions. RNA quality was assessed by spectrophotometry (A260/A280 ratio >1.8). First-strand cDNA synthesis was performed using oligo(dT) primers and reverse transcriptase. Quantitative PCR was performed in technical triplicates using SYBR Green detection chemistry with gene-specific primer sets for COL1A1 (206 bp product), LOX4 (237 bp product), and GAPDH housekeeping gene (132 bp product). Cycling conditions included initial denaturation at 95°C followed by 40 cycles of amplification. Ct values were determined using automated threshold detection. Wild-type TOC master cell line and no-template controls were included as negative controls.

#### Data analysis

Relative expression was calculated using the 2^(-ΔΔCt) method with GAPDH normalization. Stable cells without doxycycline induction served as the calibrator sample (ΔΔCt = 0). Data are presented as mean ± s.e.m. from three biological replicates. Statistical significance between induced and uninduced conditions was assessed using two-tailed Student’s t-test, with P < 0.001 considered statistically significant.

### Statistical analysis and data visualization

All statistical analyses used Python (v3.11) with scipy.stats. For multiple comparisons, Bonferroni correction was applied where appropriate. Data are presented as mean ± s.d. or mean ± s.e.m. as indicated in figure legends. Sample sizes (n) represent biological replicates and are specified in figure legends. Figures were generated using matplotlib (v3.8.2) and seaborn (v0.13.0) with publication-quality settings (300 dpi, vector graphics where applicable).

